# Proteomic changes associated with the initiation and termination of aestivation in the cabbage stem flea beetle

**DOI:** 10.1101/2025.05.21.655265

**Authors:** Gözde Güney, Doga Cedden, Stefan Scholten, Michael Rostás

**Affiliations:** Agricultural Entomology, Department of Crop Sciences, University of Göttingen, Göttingen, Germany; Department of Evolutionary Developmental Genetics, Johann-Friedrich-Blumenbach Institute, GZMB, University of Göttingen, Göttingen, Germany; Division of Crop Plant Genetics, Department of Crop Sciences, University of Göttingen, Göttingen, Germany

**Keywords:** Proteomics, summer diapause, *Psylliodes chrysocephala*, metabolism, aestivation, proteolysis

## Abstract

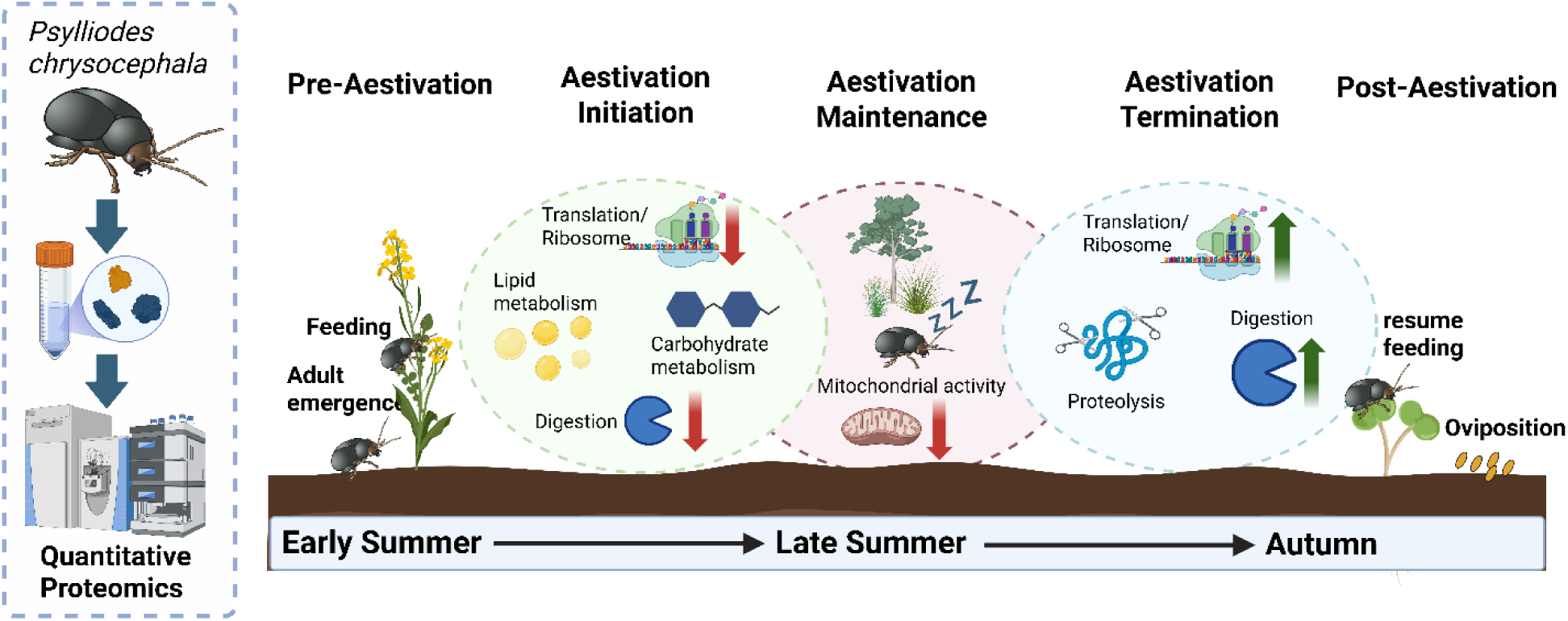

**Highlights:**

- Aestivation initiation is associated with a reduction in translation proteins
- Aestivation termination involves changes in proteolysis-related proteins
- 23-30% of the changes at the protein level were coherent with the RNA-seq study
- TAG and chitin peak, while ATP and glucose get depleted during aestivation

The cabbage stem flea beetle (CSFB, *Psylliodes chrysocephala*) is a major pest of oilseed rape crops and exhibits obligatory adult aestivation, summer diapause, that coincides with the summer season. The aestivation in CSFB is characterized by metabolic suppression, cessation of feeding, and reproductive activities. Previous investigations have employed RNA-seq to explore gene expression changes associated with aestivation, providing initial insights into the molecular pathways. However, studies assessing proteomic changes during aestivation in this species and insects more broadly have been lacking. In this study, we conducted a comprehensive quantitative proteomic analysis of adult CSFB at four time points: pre-aestivation (day 5), aestivation initiation (day 15), aestivation maintenance (day 30), and post-aestivation (day 55), to investigate proteomic changes associated with aestivation. We found that proteins related to the central dogma decreased in abundance, and metabolism-related proteins were altered during the initiation of aestivation. The proteomic changes during aestivation were minor and included reduction in mitochondrial proteins. Notably, the proteolysis-related proteins were enriched at the termination of aestivation. Interestingly, we observed discrepancies between our previous RNA-seq results and the proteomic data, particularly in the genes that increased in abundance during aestivation compared to post-aestivation. This highlights the importance of proteomic analysis for a more complete understanding of molecular mechanisms underlying aestivation. Body composition measurements showed that triglyceride and chitin levels peaked, while ATP and glucose were depleted during aestivation, following the changes in proteins belonging to different biological pathways.

## Introduction

Diapause is an alternative developmental pathway characterized by a hormonally regulated pause in development and reproductive maturation, enhanced stress tolerance, and suppression of metabolism and behavioral activity in insects (Denlinger, 2022; Gill et al., 2017). This dormant state is initiated in anticipation of unfavorable environmental conditions rather than directly responding to them, distinguishing diapause from quiescence (Radzikowski, 2013). Diapause can be classified as either facultative, triggered by environmental cues such as declining photoperiod, or obligate, where the arrest occurs at a specific life stage regardless of environmental conditions (Denlinger, 2002; Gill et al., 2017). Insects enter a preparatory phase before diapause, accumulating energy reserves and adopting protective behaviors such as burying into the soil. The accumulated energy reserves are mainly lipids and carbohydrates (Cedden et al., 2024b; Hahn and Denlinger, 2007). They are critical for survival during the dormant period and fueling post-diapause processes like migration or reproduction (Lehmann et al., 2016). While most insect diapause research has focused on winter diapause due to its prevalence in temperate regions (Tougeron, 2019), summer diapause, also known as aestivation, plays a crucial role in the survival of species that encounter seasonal aridity or heat (Saulich and Musolin, 2017).

The cabbage stem flea beetle (CSFB*, Psylliodes chrysocephala,* Coleoptera: Chrysomelidae) is a major pest of oilseed rape across northern Europe (Ewing et al., 2024; Ortega-Ramos et al., 2022). Adults damage crops by feeding on leaves, while larvae tunnel through stems and petioles, leading to growth suppression and yield loss (Alford, 1979). In the early summer, the newly emerged adults are in a stage referred to as pre-aestivation, during which they voraciously feed to accumulate energy reserves (Güney et al., 2024). During late summer, the adults obligatorily aestivate for 1-2 months, during which metabolism is suppressed, feeding activity ceases, and reproductive maturity is delayed (Güney et al., 2024; Sáringer, 1984). The aestivation is terminated in autumn, and subsequently, beetles migrate to newly sown fields to feed and oviposit (Bonnemaison & Jourdheuil, 1954; Sáringer, 1984). A previous RNA-seq study in CSFB showed that these physiological changes associated with aestivation are supported by changes in the abundance of related transcripts, such as those involved in metabolism and mitochondrial activity (Güney et al., 2024).

Management of CSFB has become increasingly difficult due to the ban on neonicotinoid seed treatments in Europe and resistance to pyrethroids (Højland et al., 2015). RNA interference (RNAi) has recently been explored as an alternative control strategy (Cedden et al., 2024a, 2023). Targeting specific molecular pathways, such as the proteasome, was able to cause up to 100% mortality within 8 days (Cedden et al., 2025, 2024a). Further studies at the molecular level might be beneficial for the development of such alternative pest control strategies against this pest, since they have the potential to reveal targets that play essential roles in the biology of the pest.

Although numerous transcriptomic studies have elucidated differentially expressed genes during diapause across various insect taxa (Güney et al., 2024; Lecheta et al., 2025; Poupardin et al., 2015; Pruisscher et al., 2022; Torson et al., 2023), much less is known about the corresponding proteomic changes. There are various translational and post-translational regulatory mechanisms described in animals, such as microRNA-mediated translational inhibition (Chekulaeva and Filipowicz, 2009) and ubiquitination-mediated degradation (Ford et al., 2024) that may uncouple protein levels from mRNA transcript levels. Hence, proteomic studies may be important to understand the molecular mechanisms underlying diapause better, as the proteome necessary for a successful diapause might not be fully captured by studying the transcriptome.

In this study, we performed quantitative proteomics in CSFB adults at pre-aestivation, early aestivation, aestivation maintenance, and post-aestivation to understand the pathways involved in aestivation. Using our previous mRNA-seq data, we evaluated similarities and discrepancies between proteomics and transcriptomics approaches in the study of aestivation. Next, we performed body composition analysis at multiple time points in CSFB adults to reveal the temporal changes associated with aestivation in energy stores and other molecules hypothesized to be altered based on proteomics analysis.

## Materials and Methods

### Insects

A laboratory colony of CSFB was maintained at 21 ± 1°C with 65 ± 10% relative humidity, following a 16:8-hour light-to-dark cycle. The insects were reared on winter oilseed rape plants at BBCH growth stages 30–35, housed in controlled rearing chambers. Newly emerged adults were collected from the soil daily and either used in bioassays or returned to the colony for breeding. To preserve genetic diversity, larvae collected annually from an untreated experimental oilseed rape field in Göttingen, Germany (coordinates: 51.564065, 9.948447) were introduced into the colony.

### Proteomics

Whole bodies of four female CSFB were collected at 5, 15, or 30 days post-adult emergence, flash-frozen in liquid nitrogen, and homogenized. Proteins were extracted using a urea denaturation buffer (6 M urea, 2 M thiourea, 10 mM HEPES, pH 8.0) at 1 ml per 100 mg of tissue. After centrifugation at 12,000 g for 5 min, protein concentration in the supernatant was measured via bicinchoninic acid assay (BCA). Proteins were precipitated using chloroform/methanol extraction, resuspended in 0.1% Rapigest SF (Waters, Yvelines, France), and digested with trypsin (1:20, Serva, Heidelberg, Germany). Following digestion, samples were acidified with trifluoroacetic acid (1:10, Sigma-Aldrich, St. Louis, MO, USA), incubated at 37 °C for 45 min, and dried using a SpeedVac. Peptides were purified using C18 stage tips. Proteomics analysis was performed with five biological replicates.

LC-MS analysis was conducted using an Ultimate 3000 system coupled to a Q Exactive HF mass spectrometer (Thermo Fisher Scientific). Peptides were separated by reverse-phase chromatography and ionized by nano-electrospray (nESI) using the Nanospray Flex Ion Source. Full MS scans were acquired in the 300–1650 m/z range at 30,000 resolution, followed by data-dependent top-10 HCD fragmentation at 15,000 resolution with dynamic exclusion. XCalibur 4.0 was used for method setup and data acquisition. MaxQuant (v2.6) and Perseus (v1.6) were used for data analysis, with LFQ values calculated using a custom CSFB protein sequence database derived from transcriptome data (Güney et al., 2024). Proteins with fewer than three peptide counts were excluded. Significance was assigned to proteins with adjusted P < 0.05 and log2 fold change > 1 or < −1. Sklearn and Seaborn were used to plot the Principal Component Analysis (PCA) graph using the LFQ values. The R package “clusterProfiler” (v4.10) was used for gene ontology (GO) enrichment analysis. We used our previous RNA-seq data (Güney et al., 2024) to assess whether the transcripts corresponding to the proteins showing differential abundance in comparing aestivation maintenance and post-aestivation increased or decreased in abundance.

### Body composition analysis

Whole body samples (n = 5-8 females) were frozen using liquid nitrogen and homogenized by grinding using a porcelain mortar and pestle. Trehalose content was quantified using 20 μL of each sample with the Trehalose Assay Kit (Megazyme, Bray, Ireland), following the manufacturer’s protocol. Glucose content was determined from 10 μL of homogenate using the D-Glucose HK Assay Kit (Megazyme). Glycogen levels were measured by incubating 20 μL of homogenate with 80 μL of 50 mM sodium acetate buffer containing 0.5 units of amyloglucosidase (TCI, Tokyo, Japan) at 37 °C for 4 h. Afterwards, 10 μL was used for total glucose quantification using the same glucose assay kit. Glycogen content was calculated by subtracting free glucose values (from untreated samples) from total glucose measured after enzyme treatment. The ATP content was measured using the ATP Assay Kit (Sigma-Aldrich).

For triglyceride (TAG) measurements, 2 μL of NP40 was added to 20 μL of homogenate, followed by centrifugation at 10,000 g for 10 min at 4 °C. Free glycerol was measured using the Free Glycerol Reagent Kit (Sigma), and total glycerol was determined with the Triglyceride Colorimetric Assay Kit (Cayman, Ann Arbor, MI, USA). TAG content was calculated as the difference between total and free glycerol measurements. Each measurement used 6–10 biological and two technical replicates on a μQuant Universal Microplate Spectrophotometer (Bio-Tek Instruments, Bad Friedrichshall, Germany).

The chitin content was quantified using a modified method derived from a previous study (Lehmann and White, 1975; Zhang and Zhu, 2006). Briefly, each insect was homogenized in distilled water with metallic beads, and the homogenate was centrifuged (1800 g for 15 min at room temperature). The pellet was washed and resuspended in 0.2 ml of 3% SDS (sodium dodecyl sulfate, Sigma-Aldrich). Following heat treatment (100 °C for 15 min) and additional centrifugation (10 min, 1800 g), the pellet was washed with 0.2 ml of distilled water and then resuspended in 0.2 ml of 14 M KOH at 130 °C for 1 h to deacetylate the chitin. Ice-cold 0.4 ml of 75% ethanol was then added to the samples, which were further incubated on ice for 15 min. A 30 μL suspension of Celite 545 (Sigma-Aldrich) was added to the samples and centrifuged (1800 g at 4 °C for 5 min). The chitosan pellet was washed with 0.2 ml of 40% cold ethanol and then with 0.2 ml of distilled water. The pellet was resuspended in 0.2 ml of distilled water by vortexing. For the colorimetric assay, 100 µL of the chitosan solution was sequentially treated with 50 µL of 10% NaNO₂ and 50 µL of 10% KHSO₄, mixing three times during a 15-minute incubation at room temperature. Following centrifugation for 5 minutes at 1800 g at 4 °C, 60 µL of supernatant was transferred to a microcentrifuge tube, and 20 µL of NH₄SO₃NH₂ was added and shaken for 5 minutes at room temperature. After adding 20 μL of freshly prepared MBTH reagent (3-methyl-2-benzothiazolone hydrazone hydrochloride hydrate, TCI, Tokyo, Japan), the mixtures were incubated at 100 °C for 5 minutes, cooled at room temperature, and then 100 μL of the sample was transferred to a 96-well plate. Absorbance was measured at 650 nm using a microplate reader. Finally, chitin content was expressed in glucosamine equivalents according to a calibration curve created from the measurement of known concentrations of glucosamine (Sigma-Aldrich).

The body composition measurements were analyzed using one-way ANOVA, followed by Tukey’s multiple comparisons test to compare different time points with each other. P values below 0.05 were deemed statistically significant.

## Results

### Differentially abundant proteins in aestivation

We performed quantitative proteomics on whole bodies of CSFB at 5, 15, 30, and 55 days post-emergence, corresponding to pre-aestivation, early aestivation, aestivation maintenance, and post-aestivation, respectively. In total, 2,216 proteins were quantifiable in at least one of the stages. Principal component (PC) analysis revealed clear clustering of the different stages (PC1: 29.5%, PC2: 11.6%, n = 5, Fig. 1A). The most distinct cluster was formed by the aestivation maintenance (30-day-old) adults, which were especially distinguishable from pre-aestivation adults along PC1. Interestingly, the confidence intervals of the early aestivation and post-aestivation clusters partially overlapped.

**Figure 1.**
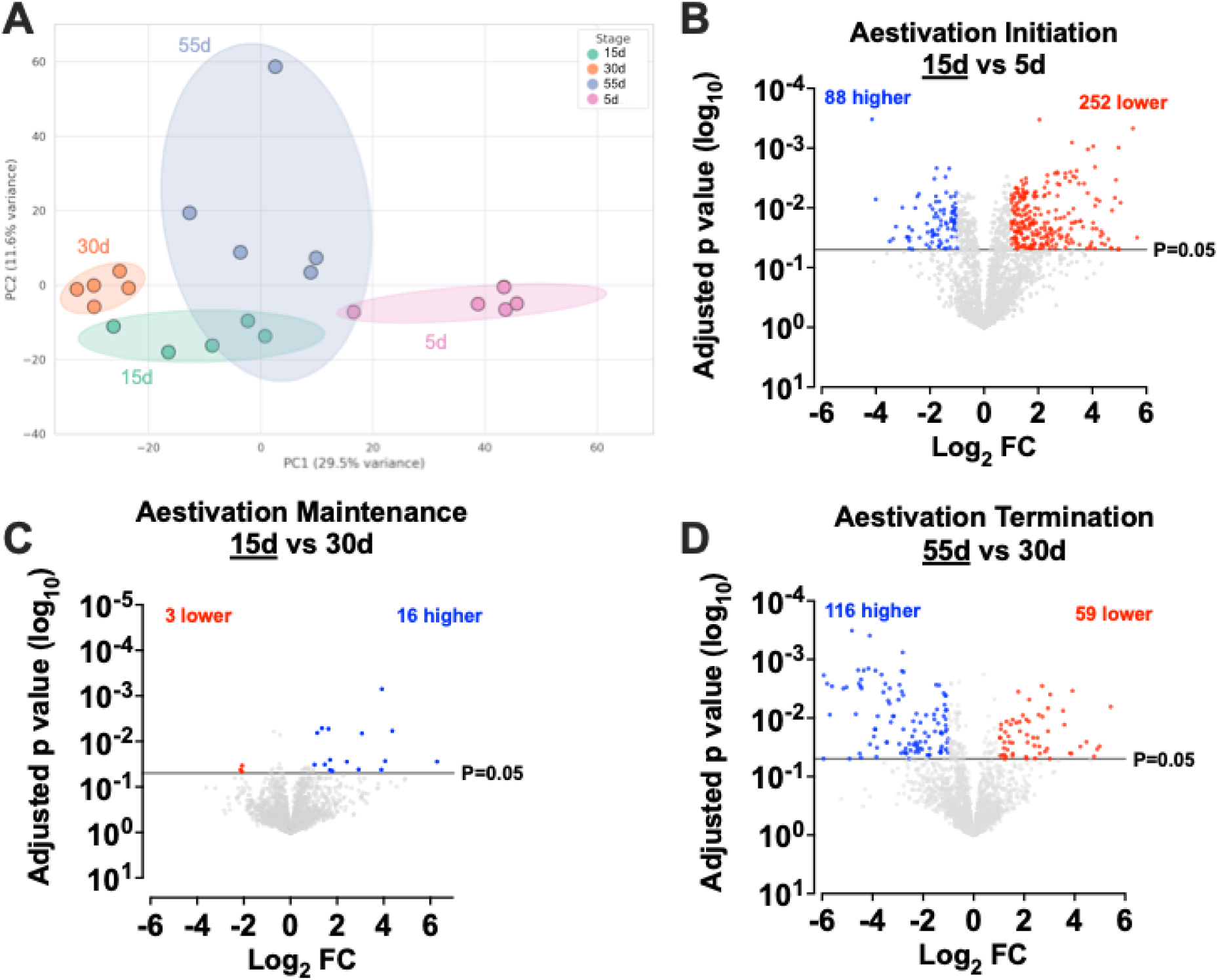
Differentially abundant proteins at different life stages in the cabbage stem flea beetle. A) Principal component analysis (PCA) of protein abundances in 5-, 15-, 30-, and 55-day-old adults, representing pre-aestivation, early aestivation, aestivation maintenance, and post-aestivation. B-D) Volcano plots showing the Log_2_ fold change (FC) in LFQ values and corresponding Log_10_ adjusted P values in the comparison of 15-vs. 5-day-old adults (Aestivation initiation, Panel B), 15-vs. 30-day-old adults (Aestivation maintenance, Panel C), 55-vs. 30-day-old adults (Aestivation termination, Panel D). Blue dots indicate proteins with significantly higher abundance, while red dots indicate those with significantly lower abundance in the underlined stage. Proteins with adjusted P values below 0.05 and Fold changes higher or lower than 1 and −1 were considered to be significantly changed in abundance. Of note, the results presented in Panel B were also previously used as a control experiment in a previous study (Güney et al., 2025b).

Next, we performed three pairwise comparisons to assess changes during aestivation initiation, maintenance, and termination. During aestivation initiation, 252 proteins showed lower abundance. In contrast, only 88 proteins had higher abundance in early aestivation compared to pre-aestivation (Fig. 1B). Aestivation maintenance involved 16 proteins with reduced abundance and only 3 proteins with increased abundance compared to the pre-aestivation stage (Fig. 1C), indicating a further reduction in protein abundance as aestivation progresses. During aestivation termination, 116 proteins exhibited higher abundance, while 59 proteins were less abundant in the post-aestivation stage (Fig. 1D) compared to aestivation maintenance. We also observed that both early aestivation and aestivation maintenance beetles had lower total protein abundance compared to pre-aestivation beetles (Fig. S1, P < 0.05 by Tukey’s test, n = 5). Overall, these results suggested that most proteins were reduced in abundance during aestivation, whereas the termination of aestivation is associated with a rebound in protein levels.

### Enriched pathways in aestivation

To gain biological insights into the initiation, maintenance, and termination of aestivation, we performed gene ontology (GO) enrichment analyses on proteins showing significant differences in abundance.

In the early phase of aestivation (15 vs. 5 days), proteins decreasing in abundance were primarily associated with ribosome-related and other translation-related processes (Fig. 2A). Additional enriched terms included those related to metabolism, such as carbohydrate metabolic process, digestion, and fatty acid biosynthesis. In contrast, proteins with higher abundance in early aestivation (5 vs. 15 days) included those involved in metabolism, such as the tricarboxylic acid (TCA) cycle, as well as processes linked to muscle tissue, including myofibril assembly and muscle-tendon junction organization (Fig. 2B).

**Figure 2.**
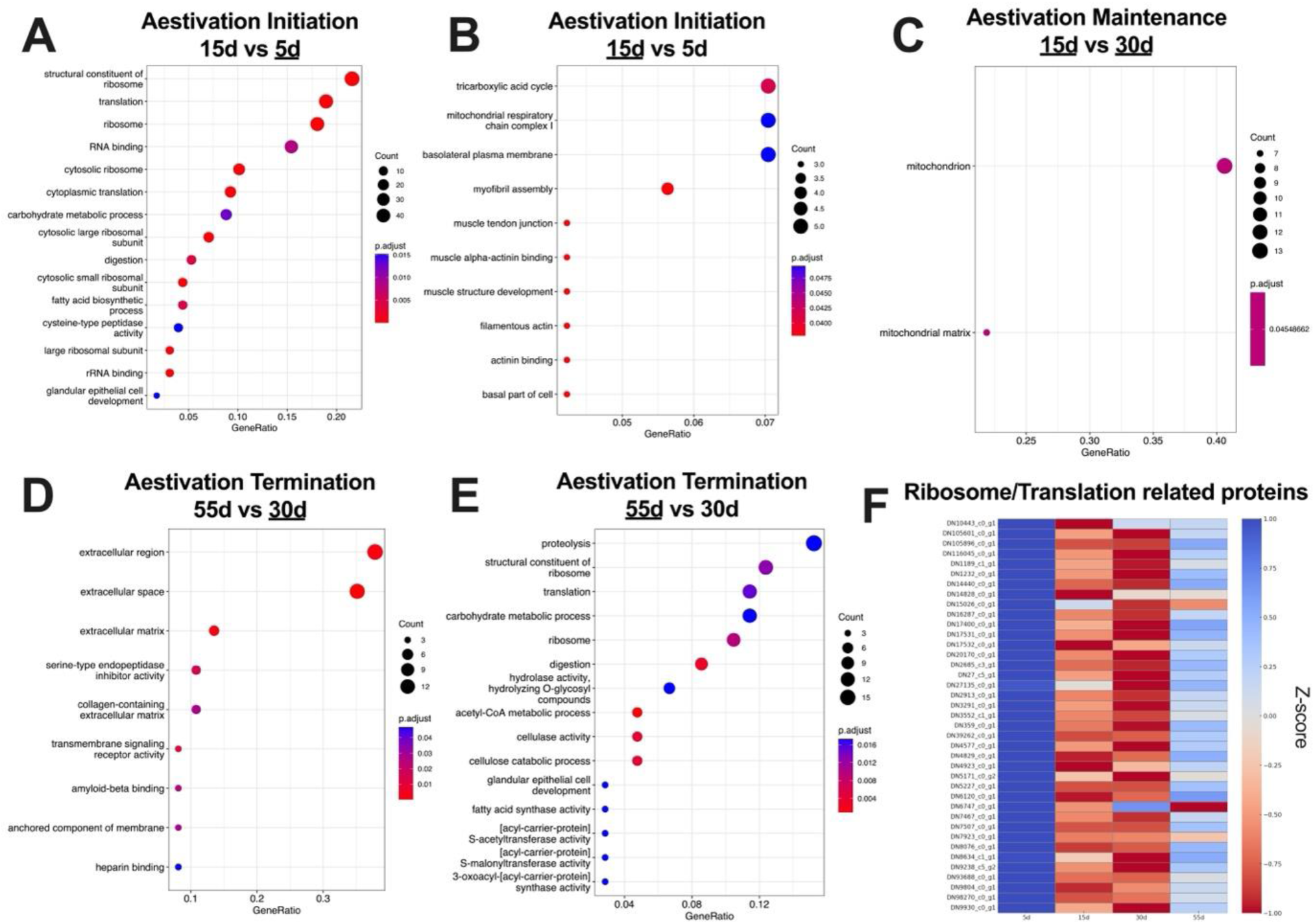
Gene Ontology (GO) enrichment analysis of differentially abundant proteins across life stages of the cabbage stem flea beetle. GO term enrichment was performed on proteins that were significantly higher or lower in abundance in the stage comparisons shown in Figure 1. The underlining indicates the stage at which the proteins had significantly higher abundance, except for Panel C, where all differentially abundant proteins were combined because separated data sets did not reveal any enrichment. A-B) GO enrichment in proteins with higher abundance during aestivation initiation (15 vs. 5 days). C) GO enrichment in proteins associated with aestivation maintenance (30 vs. 15 days). D-E) GO enrichment in proteins associated with aestivation termination (55 vs. 30 days). Only GO terms with adjusted p-values < 0.05 are shown together with the count of protein hits associated with the term. Gene ratio indicates the ratio of protein hits over the total number of proteins associated with the respective GO term.

During the maintenance phase of aestivation (30 vs. 15 days), GO enrichment analysis revealed minimal changes. Only two mitochondrial-related terms passed the significance threshold (Fig. 2C). As the majority of proteins showed lower abundance in this phase compared to early aestivation, these findings suggest repression of metabolism as aestivation progresses.

At the termination of aestivation, proteins with higher abundance in the aestivation compared to post-aestivation (30 vs. 55 days) were primarily associated with extracellular structures, including the extracellular region and extracellular matrix (Fig. 3D). The termination of aestivation (55 vs. 30 days) was marked by an increase in the abundance of translation and proteolysis-related proteins in post-aestivation, suggesting an extensive remodeling of the proteome (Fig. 3E).

**Figure 3.**
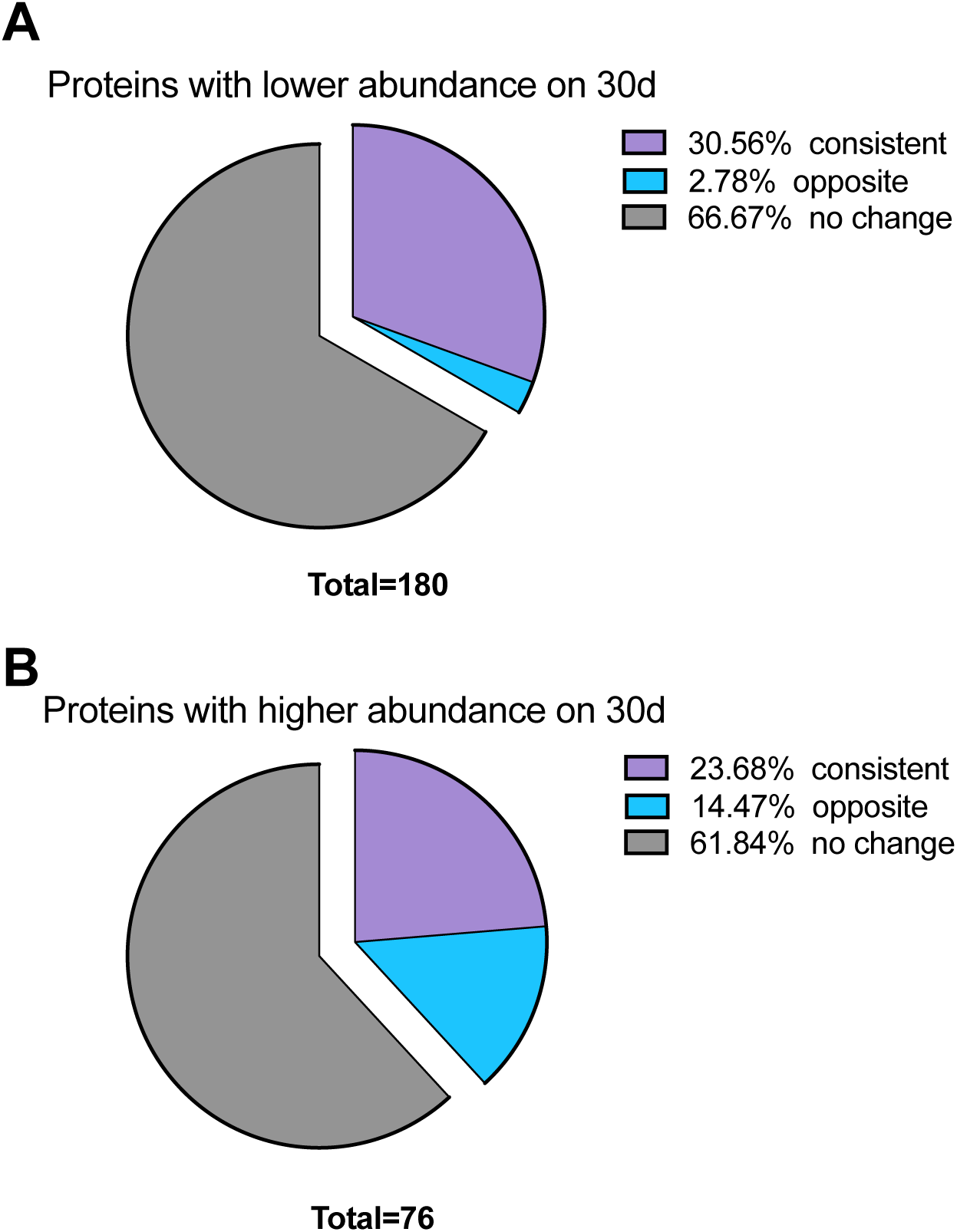
Comparison of proteomics and transcriptomics results in the study of aestivation termination. A) Proportions of proteins with lower abundance on 30-day-old (aestivation maintenance) CSFB that showed consistent, opposite, or no change at the transcript level. B) Proportions of proteins with higher abundance on 30-day-old CSFB that showed consistent, opposite, or no change at the transcript level. The transcriptome data were taken from a previous study for comparison (Güney et al., 2024).

The heatmap (Fig. 2F) illustrates the Z-score normalized abundance of proteins associated with the ribosome and other translation-related processes that showed a significant difference in at least one pairwise comparison. The pattern indicates that the pre-aestivation stage is the most active in terms of protein synthesis. This is followed by a notable reduction in the abundance of these proteins during early aestivation and maintenance. A subsequent increase is observed during the termination phase, although protein levels remain slightly below those seen in the pre-aestivation stage.

### Comparing proteomics with transcriptomics

Next, we sought to understand how far our previous transcriptomics study reflected the proteomic changes identified in this study. For this, we selected the aestivation termination as the study case, as we performed transcriptomics and proteomics analyses using adults of the same ages (30- and 55-day-old adults). One limitation of this comparison is that not all proteins detectable at the transcript level are quantifiable via proteomics. Since all quantifiable proteomics were also quantifiable through transcriptomics in our previous study, we focused on the differentially abundant proteins and checked whether the respective transcripts showed differential expression as well. Also, we considered all proteins below the adjusted P value of 0.05 to be differentially abundant for this comparison.

Applying this analysis to the proteins with lower abundance in diapause maintenance compared to post-aestivation revealed that 30.56% of these proteins showed consistent differential abundance at the transcript level (i.e., lower transcript abundance, Fig. 3A). The Majority of proteins (66.67%) showed no differential expression at the transcript level. A relatively small proportion of these proteins (2.78%) showed opposite regulation at the transcript level (i.e., higher transcript abundance).

Next, we performed the same analysis for the proteins with higher abundance in aestivation maintenance compared to post-aestivation (Fig. 3B). In this analysis, 23.68% of these proteins showed consistent differential abundance at the transcript level (i.e., higher transcript abundance). Interestingly, 14.47% of these proteins showed the opposite relationship, where the respective transcript had lower abundance during aestivation maintenance compared to post-aestivation. Similar to the case above, the majority of these proteins (61.84%) showed no differential expression at the transcript level. Overall, these results suggest that there are notable discrepancies between changes at the protein and corresponding transcript levels, and this was especially prominent when we compared proteins with higher abundance in aestivation maintenance.

### Body composition analysis

The enrichment analyses on differentially abundant proteins suggested that aestivation involves changes in metabolism and extracellular space-related activities. Here, we performed body composition analysis in adults at multiple time points. Triglyceride (TAG) levels indicated rapid accumulation of TAG reserves during pre-aestivation and stable levels during early aestivation (Fig. 4A). In contrast, TAG levels were reduced between 30- and 36-days post-emergence, suggesting the utilization of TAG reserves during late aestivation.

**Figure 4.**
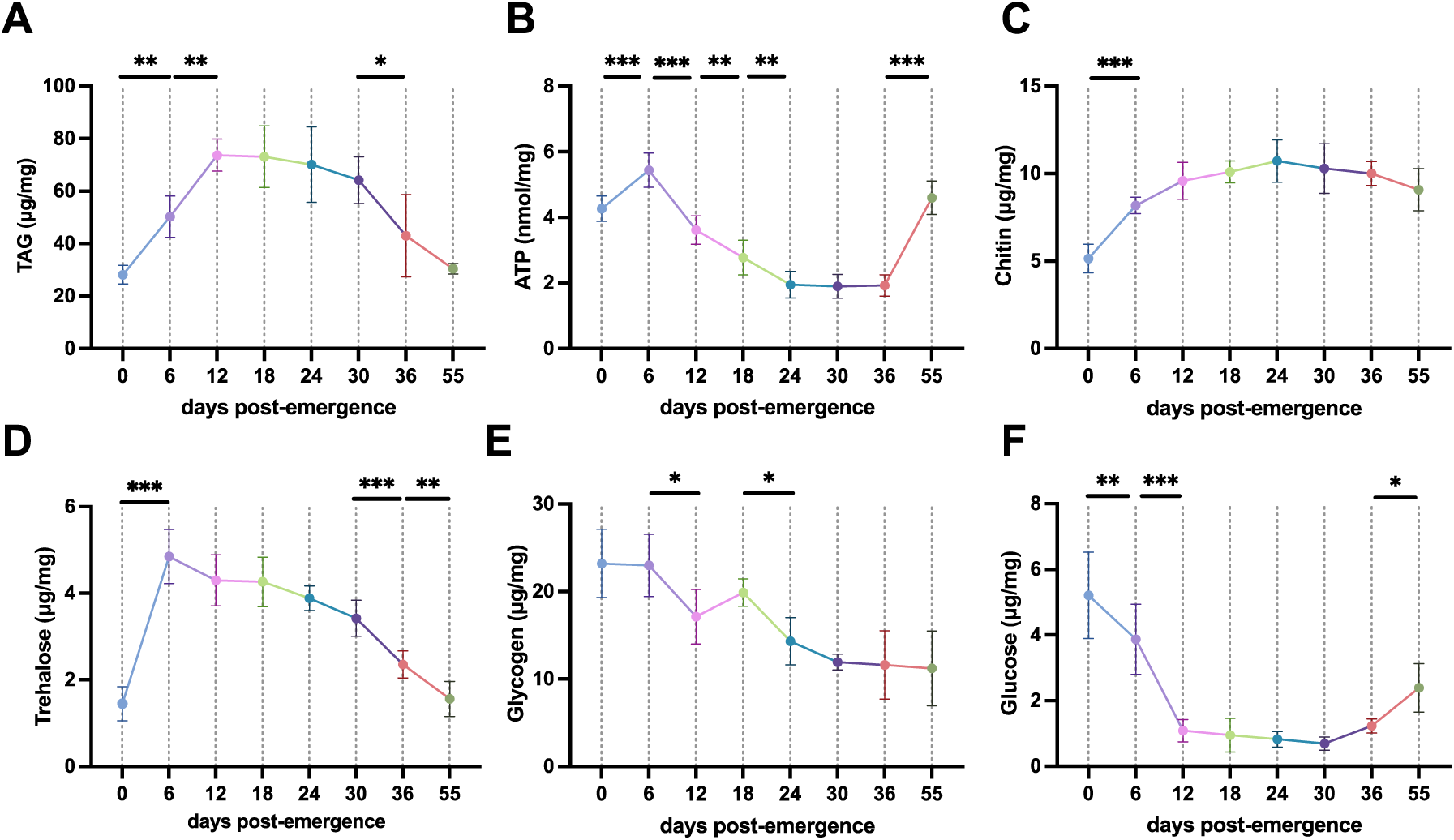
Temporal body composition analysis in adult cabbage stem flea beetles. Whole bodies of females at different time points following adult emergence were used to measure triglyceride (TAG, A), adenosine triphosphate (ATP, B), chitin (C), trehalose (D), glycogen (E), and glucose (F) contents (n = 5–8). Days 0-12 represent pre-aestivation, 18-36 represent aestivation, and day 55 represents post-aestivation. One-way ANOVA and Tukey’s multiple comparison tests assessed statistical differences. Statistical significance is indicated as follows: P < 0.05 (*), P < 0.01 (**), and P < 0.001 (***).

ATP levels peaked shortly after emergence at 6 days post-emergence but were notably reduced throughout aestivation, only to rebound during post-aestivation (Fig. 4B). Chitin levels showed a significant increase during pre-aestivation at 6 days post-emergence (P < 0.05), likely as part of the regular adult emergence process (Fig. 4C). The peak in chitin content was observed during aestivation maintenance, albeit the increase was very gradual and not associated with significant changes between adjacent time points (P > 0.05).

Trehalose levels showed a prominent peak in pre-aestivation (day 6) and gradually declined during aestivation and post-aestivation (Fig. 4D). Glycogen levels were highest in newly emerged adults and then gradually declined during aestivation, with no recovery in post-aestivation (Fig. 4E). Glucose levels were also highest in newly emerged adults but were dramatically reduced during pre-aestivation (day 6) and aestivation (day 30, Fig. 4F). In contrast to glycogen levels, glucose levels showed a slight recovery during post-aestivation (day 55). Overall, the body composition analysis revealed dynamic changes before, during, and after aestivation, highlighting that while certain stores, such as TAG, are retained, others, like glucose, get depleted during aestivation.

## Discussion

The CSFB represents a valuable insect model for studying aestivation due to the availability of various molecular tools established in this species, including RNA-seq and RNAi (Cedden et al., 2024a; Güney et al., 2024). Moreover, unlike the majority of insect diapause studies that focus on facultative diapause, the obligatory nature of aestivation in CSFB eliminates the need for external environmental manipulations to induce diapause, thus reducing experimental confounders and allowing for more precise molecular investigations. Here, we used aestivating CSFB as a model to study diapause-associated changes using quantitative proteomics, which has not been incorporated into diapause research as much as RNA-seq.

Our proteomic analysis revealed that a prominent change associated with aestivation was a decrease in the abundance of most proteins compared to non-aestivation stages. The GO enrichment analysis revealed that many proteins with lower abundance in aestivation were involved in the central dogma and protein synthesis, further supporting that aestivation is associated with an overall suppression of cellular activities that are maintained through protein homeostasis. A general reduction in protein levels during diapause has been previously shown in various insects, including *Culex pipiens* (Zhang et al., 2019) and *Bombus terrestris* (Colgan et al., 2019). Similarly, a study focusing on aestivation in the grain chinch bug, *Macchiademus diplopterus* found a decrease in the number of proteins as the insects were transitioning into aestivation, suggesting the hypometabolic state is ensured through changes in protein levels (Smit et al., 2024).

Interestingly, we observed a rebound at the aestivation termination in the abundance of most proteins that had decreased in abundance at the aestivation initiation. This finding aligns with the physiological requirements of post-aestivation adults, who must resume feeding and prepare for reproduction and dispersal (Güney et al., 2024; Sáringer, 1984). Similarly, the study in *M. diplopterus* also found a rebound in the protein abundances towards the termination of aestivation (Smit et al., 2024). However, in the current study, we observed that most translation-related proteins that decreased during aestivation did not return to their pre-aestivation levels. This raises the idea that pre-aestivation might be a more cellularly demanding stage compared to post-aestivation, despite the latter being associated with reproductive activity. A similar conclusion was also made in our previous study comparing the metabolic rates and transcriptomes of pre- and post-aestivation stages in CSFB (Güney et al., 2024). One potential explanation is that pre-aestivation is cellularly very active because it needs to synthesize metabolites and other molecules necessary for the subsequent aestivation stage.

In contrast, we found that proteomic changes observed during the maintenance phase were minimal, with only a few proteins showing differential abundance, likely further repressing the metabolism. This suggests that once the diapause program is established, CSFB’s cellular physiology remains relatively stable until diapause is terminated, as observed in other insects (Lehmann et al., 2020).

Body composition analysis suggested that TAG, chitin, and trehalose levels seem to be increasing during pre-aestivation in anticipation of aestivation. Previous studies showed that TAG and trehalose play important roles in diapause by acting as energy stores (both) or resilience against abiotic stress (latter) (MacRae, 2010; Wang et al., 2020; Zhang et al., 2025). The accumulation of TAG reserves was in line with proteomic indications of altered lipid metabolism at the aestivation initiation, revealed by the GO enrichment analysis. The reduction in ATP level throughout aestivation and its recovery in post-aestivation further reflect suppressed and reactivated metabolic activity, respectively, which was also observed in our previous study in CSFB, where the CO_2_ output showed the same pattern (Güney et al., 2024) and other insects such as the Colorado potato beetle, *Leptinotarsa decemlineata* (Lehmann et al., 2014). A recent study in CSFB investigated trehalose dynamics in further detail in aestivation and found that trehalose transporters are involved in intricate regulation of trehalose in specific tissues such as the fat body, and play a role in the heat resilience in aestivating beetles (Güney et al., 2025a).

It remains unclear whether the increase in chitin levels during pre-aestivation has functional relevance for aestivation or simply coincides with post-eclosion maturation. If an increase in chitin level has a functional role in aestivation, it could be related to the fortification of the cuticle (Khajuria et al., 2013), which might explain the higher heat stress tolerance observed in aestivating CSFB (Güney et al., 2024). Functional studies using RNAi to target chitin metabolism-related genes might elucidate whether chitin accumulation is playing a role in aestivation.

Our comparison of proteomic and transcriptomic data revealed notable discrepancies between mRNA levels and protein abundances at the termination of aestivation. A significant proportion of proteins showing increased abundance in post-aestivation adults were not accompanied by similar increases at the transcript level. In fact, for proteins with higher abundance during aestivation maintenance, a notable subset (14.47%) exhibited lower transcript levels at the same time point. These observations underscore the critical role of regulation at the translational and post-translational levels in diapause, which may include proteasome-mediated degradation or microRNA-mediated translational inhibition previously investigated in CSFB (Cedden et al., 2025; Güney et al., 2025b). The decoupling of transcription and translation highlights the complexity of regulatory networks underlying diapause and reinforces the necessity of proteomic analyses to complement transcriptomics in diapause research. A potential reason for such decoupling could be the rapid and independent evolution of diapause in insects, which may have led to the establishment of complex gene regulatory mechanisms governing diapause (Batz et al., 2020; Powell et al., 2020; Ragland et al., 2019).

Despite discrepancies at the single gene level, functional enrichment analyses revealed overlapping biological themes across both datasets. For instance, terms related to digestion, carbohydrate metabolism, and lipid biosynthesis were consistently associated with aestivation in both the proteomics approach in this study and the transcriptomic approach in our previous study (Güney et al., 2024). This convergence at the pathway level indicates that both proteomic and transcriptomic approaches provide similar insights into the broader physiological processes regulated during diapause. Interestingly, one term, proteolysis at aestivation termination, was not identified in our previous transcriptomics study. This finding suggests that the termination of aestivation involves the enzymatic breakdown of proteins that are no longer needed, potentially providing amino acids for the synthesis of a metabolically active proteome during the post-aestivation. Proteolysis activity has been previously implicated in the diapause of other insects, including *Helicoverpa armigera* and *Ostrinia furnacalis* (Su et al., 2024; Yang et al., 2024).

### Conclusions

This study presents a comprehensive proteomic analysis of obligatory aestivation (i.e., summer diapause) in the cabbage stem flea beetle, revealing molecular changes during the initiation, maintenance, and termination of diapause. Aestivation was marked by an overall reduction in protein abundance, enabled by the reduction in translation-related proteins. The proteome remained relatively stable during aestivation but was followed by a partial rebound upon termination. Comparing the proteomics results with transcriptomic data for the study of aestivation termination showed that most changes at the protein level are not reflected at the transcript level, highlighting the likely importance of translational and post-translational regulations in diapause. Proteomic changes were consistent with body composition measurements, including triglyceride levels, which increased during pre-aestivation and were consumed during the aestivation maintenance. These findings underscore the value of proteomics in understanding diapause at the molecular level and point to future directions, including the study of protein modification and turnover mechanisms important for diapause.

## Acknowledgments

The authors, DC and GG, were funded by the Deutscher Akademischer Austauschdienst (DAAD) research grants program. The authors thank Dr. Oliver Valerius and Dr. Kerstin Schmitt for the proteomics method.

## Author Contributions

Gözde Güney: Writing – original draft, Visualization, Validation, Methodology, Investigation, Formal analysis, Data curation, Conceptualization. Doga Cedden: Writing – review & editing, Visualization, Software, Formal analysis, Data curation, Conceptualization. Stefan Scholten: Writing – review & editing, Resources, Funding acquisition. Michael Rostás: Writing – review & editing, Resources, Funding acquisition

## Supplementary

**Figure S1.**
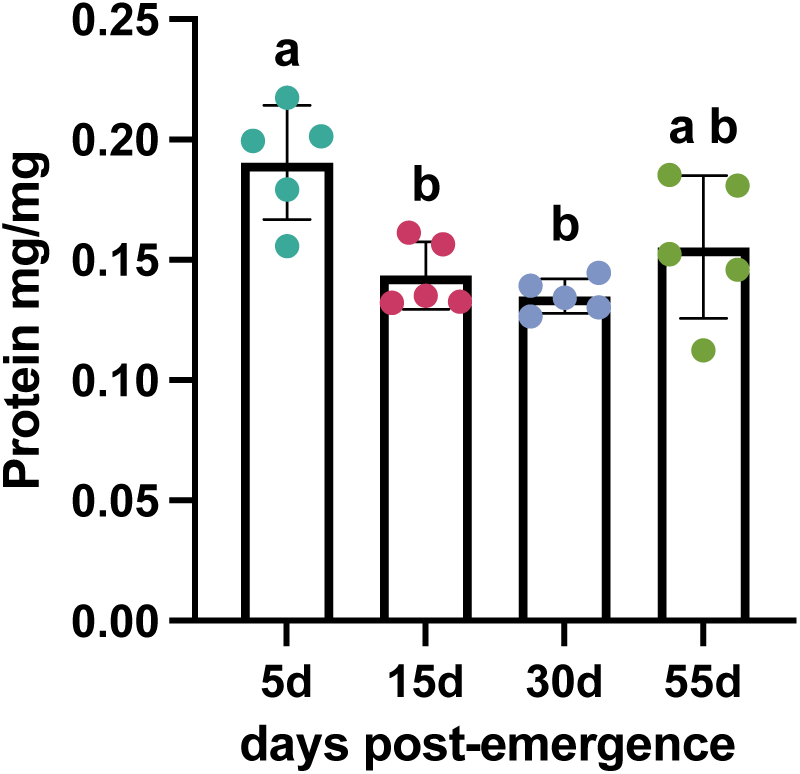
Protein contents measured in CSFB whole bodies (n = 5) at different time points. Different letters denote statistical significance as determined by Tukey’s multiple comparison test following one-way ANOVA.

